# The Medial Temporal Lobe Supports Mnemonic Discrimination For Event Duration

**DOI:** 10.1101/2020.04.13.039545

**Authors:** Nathan M. Muncy, C. Brock Kirwan

## Abstract

Time has an integral role in episodic memory and previous work has implicated the medial temporal lobe in both representing time and discriminating between similar memory traces. Here we developed a novel paradigm to test mnemonic discrimination for temporal duration, as previous temporal work has largely focused on order or maintaining information over time. Thirty-five healthy, young adults completed a continuous-recognition temporal discrimination task in which participants were tasked with detecting a change of stimulus duration on the order of 0.5 seconds and whole-brain high-resolution fMRI data were acquired during this process. Analyses of behavioral results indicate that participants were successful at detecting whether the stimulus duration changed. Further, fMRI analyses revealed that successful trial performance was associated with differential processes in the left entorhinal and perirhinal cortices. Specifically, the left entorhinal cortex was differentially engaged during encoding trials that preceded Target detection, and the left perirhinal cortex was differentially engaged during successful test phase Target and Lure detections. These findings suggest that the entorhinal cortex is involved in the encoding of temporal context information and that the perirhinal cortex is representing the conjunction of item and context during retrieval.

## 1 Introduction

Temporal information is critical for episodic memory but rather unique in the sense that the temporal properties do not directly map to any receptor in the same way that other sensory modalities do. Rather, time is a “hidden dimension” (Howard et al., 2014); the information is inferred through intermediate neural representations rather than actual interactions with the environment. Recent work suggests that it is precisely due to such intermediate representations that disparate information is able to be associated (Howard & Eichenbaum, 2013; Howard et al., 2014; MacDonald et al., 2011). Take, for example, the various information types presented to the student during a college lecture; their memory of the lecture will likely contain visual, spatial, and auditory properties as well as the temporal properties of when the lecture occurred and possibly the length of the lecture as well. In this example, the differing stimulus modalities arrive at the central nervous system with discrete but meaningful temporal associations which are then integrated into a neural representation of time, thereby allowing for the disparate information to be associated into a unique and unitized (Clewett & Davachi, 2017) memory representation.

The hippocampus receives multi-modal input from various cortical regions (Aggleton, 2012; Baldassano et al., 2016; Cabeza et al., 2008; Eichenbaum et al., 2012; Mayes et al., 2007; Qin et al., 2016; Rockland & Van Hoesen, 1999; Suzuki & Naya, 2014; Thierry et al., 2000). As such, the hippocampus is ideally situated to form associations among disparate sensory modalities, stitching them together into a single representation that will subsequently be retrievable (Backus et al., 2016; DuBrow & Davachi, 2014; Howard et al., 2014; Thavabalasingam et al., 2018). Previous work has demonstrated that the hippocampus also generates a representation of time: work by Pastalkova et al. (2008) demonstrated that the hippocampus contained a self-organized internal mechanism by which temporal information was maintained. Further, more recent modeling (Howard & Eichenbaum, 2013) has demonstrated that the hippocampus is capable of producing scale-invariant representations of time. It is through these representations that the recovery of contextual temporal information in episodic memory is possible, thereby allowing for the “backward jump in time” (Howard & Eichenbaum, 2013). MacDonald et al. (2011) has also shown that certain ensembles of hippocampal neurons, termed “time cells”, had peak activity at different and sequential periods, effectively serving to “bridge the delay period” and maintain information through a representation of time. Further work by Howard et al. (2014) argued that varying time scales may also be represented by differing neural ensembles coding for different time constants. Such time constants are then thought to be combined with unimodal input from various cortical regions, thereby forming “holistic representation that captures the relationships between stimuli separated by time and space” (Howard et al., 2014, p. 4703). As such, not only does the hippocampus serve to stitch together multiple information modalities in time, it also constructs the temporal foundation upon which such associations are formed.

Given the intimate relationship between time and information in episodic memory, it is possible that the hippocampus would be sensitive to event duration at different time scales. In addition to the varying scales formed via neural ensembles (Howard et al., 2014), the hippocampus has been shown to be sensitive to time on the order of seconds (Barnett et al., 2014; Thavabalasingam et al., 2018) as well as on the order of minutes (Montchal et al., 2019); for an excellent review, see Lee et al. [2020]]). Given that pattern separation processes have been detected within hippocampal circuitry (Gilbert & Kesner, 2006; Gilbert et al., 1998; Gilbert et al., 2001; Hunsaker & Kesner, 2013), that pattern separation processes are information agnostic (Huffman & Stark, 2014; Yassa & Stark, 2011), and hippocampal sensitivity to temporal information, we expect the hippocampus to be capable of discriminating between similar but different temporal properties within episodic memory. Many have studied temporal pattern separation or mnemonic discrimination (Gilbert et al., 2001; Hunsaker & Kesner, 2013; Montchal et al., 2019), but these paradigms have largely tested the sensitivity of hippocampal processes to order of stimuli. While the order of events is important for episodic representations, event duration is also critical. Importantly, while order contains temporal components, order itself is not inherently temporal, given that “[o]rdinal information can be extracted from a temporal representation, but the converse is not true” (Howard & Eichenbaum, 2013, p. 1212). This idea was addressed by Montchal et al. (2019) when discussing the importance of considering the relationship of sequence and duration, concluding “[i]t is likely that both types of information are important for making temporal judgments” (p. 287), and, perhaps somewhat conversely, it has recently been suggested that order or an evolving contextual representation is integral and perhaps necessary for memory of event duration (Lee et al., 2020). To date, however, only a single study has aimed at testing episodic memory for duration in humans (Thavabalasingam et al., 2019), where CA1 was demonstrated to be critical for duration memory. Related work include a set of experiments conducted by Barnett et al. (2014), which clearly demonstrated that human participants were capable of detecting second and sub-second changes to both inter-stimulus and event durations when presented with a series of images, a behavior which was associated with differential hippocampal activity. Also, work by Palombo et al. (2019) demonstrated that medial temporal lobe patients were successful at detecting whether a set of stimuli changed in duration from that of another set. We note, though, that both experiments were operating within a time frame largely governed by working memory, and Thavabalasingam (2019) is the only study of which we are aware that tested episodic memory for temporal duration. So, while the hippocampus is robustly implicated in temporal processing, episodic memory, and mnemonic discrimination, it is yet unknown whether mnemonic discrimination for the episodic memory of temporal duration is possible, and if so, what are the governing processes.

We have therefore designed a set of experiments to (1) determine whether participants can distinguish between highly similar temporal durations in episodic memory and (2) implicate the important neural regions associated with detecting changes in temporal information. We note that the paradigm employed was conceptualized as an attempt to isolate temporal duration information from order as well as test episodic memory precision for temporal duration. In the first experiment we hypothesized that participants would be (a) sensitive to changes in temporal duration on the order of 0.5 seconds, (b) differentially sensitive when fewer stimuli separate the study trial from the Target or Lure, and (c) differentially sensitive to increases than decreases in stimulus duration. In the second experiment, we hypothesized that (d) the medial temporal regions, and in particular the CA1, would be involved in successful Lure detection, given that this region is largely implicated by extant research (Azab et al., 2014; Gilbert et al., 2001; Howard et al., 2014; Kitamura et al., 2014; Montchal et al., 2019; Schapiro et al., 2016; Thavabalasingam et al., 2018; Thavabalasingam et al., 2019); we expected an increased BOLD response in the CA1 that corresponded with successful Lure detection relative to an error. Further, we expected that (e) a gradient would exist in which the anterior hippocampus (head) would show greater sensitivity to temporal information (DuBrow & Davachi, 2014; Montchal et al., 2019; Thavabalasingam et al., 2019) than more posterior regions of the hippocampus.

## 2 Methods - Behavioral Experiment 1

### 2.1 Participants

Forty healthy, young-adult participants were recruited from the Brigham Young University student population. This number was determined via a priori G*Power (Bruin, 2018) calculations, where the test family = F tests, statistical test = ANOVA: repeated measures, within factors, Effect size = 0.25, Power = 0.95, Number of groups = 1, Number of measurements = 4, Correlation = 0.5, and Nonsphericity correction = 1; such input yielded a required sample size of 36. The inclusion criteria consisted of speaking English as a native language and having normal or corrected-to-normal vision. Exclusion criteria consisted of a history of psychologic disorder or brain injury, and color blindness. The university’s Institutional Review Board approved the research, and all participants gave written informed consent prior to participation. Participants were compensated for their time via course credit.

### 2.2 Temporal Discrimination Task

The behavioral task consisted of six blocks of a continuous-recognition paradigm in which a continuous stream of trials were presented (Figure 1). Each trial consisted of a fixation cross (a black cross on a white background) displayed for 0.5s, the stimulus (a colored image of an everyday object) for a variable duration (see below), a colored masking image to combat iconic memory presented for 0.3s, and a response screen displayed for 2.5s. If the stimulus was novel (a study trial or a Foil), one of three response screens was presented at random prompting the participant to indicate a judgment about a physical property of the object via button press. These response options were a) whether the object was Smooth, Rough, or Sharp, b) whether the object was made of Metal, Plastic, or Other, and c) whether the item’s weight was Heavy, Medium, or Light. Each response option was presented an equal number of times. The aim of these response options was to maintain engagement in the task given that the perception of time is related to attention (Coull et al., 2004; Radua et al., 2014). All study trials and Foils were presented for either 1 or 1.5s. For test trials (i.e. a Target or Lure), the response screen prompted the participant to decide whether the stimulus was presented for either a Longer, Same Time, or Shorter duration than the study trial. For Target trials, stimulus and duration were identical to the study trial. Lure trials also had an identical stimulus to the study trial, but the presentation duration changed by 0.5s. Specifically, the Lure was presented for 1.5s if the study counterpart was presented for 1s, and vice versa if the study trial was presented for 1.5s. In this way all stimuli (Target, Lure, Foil) were presented for either 1 or 1.5s but Lure trials either increased or decreased by 0.5s. In total, the earliest a test trial could occur, assuming Lag = 4 (see below) and all intermediate durations = 1s, was 17.2 seconds, and the latest was 57.6 seconds (Lag = 12, Duration = 1.5s). Prior to the first block, the participant was instructed in the task via a video with explicit instructions as well as a practice period, and no feedback on performance was given during the task. Stimuli were presented in PsychoPy2 (Peirce et al., 2019).

**Figure 1:**
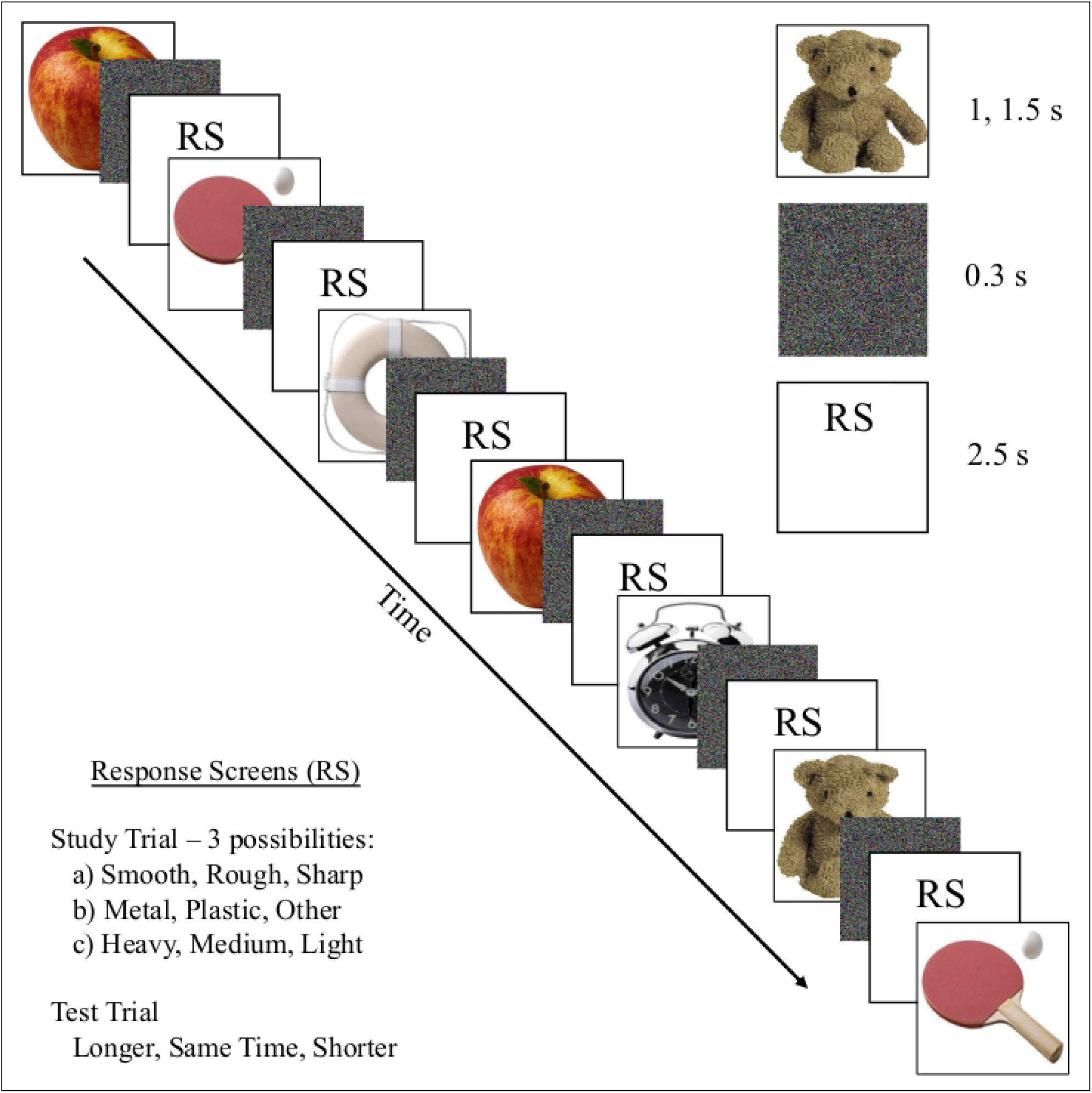
Temporal discrimination paradigm. A trial started with a fixation cross (0.5s, not shown) and then a stimulus was presented for either 1 or 1.5s. Following the stimulus, a colored mask was presented for 0.3s, and then a response screen (RS) was presented for 2.5s. Four RS were possible: for study trials one of three memory prompts for visual properties were presented, and for test trials a memory prompt for stimulus duration was presented. Targets and Lures (test trials) followed their study trial counterpart by either 4 or 12 trials. Targets had the same duration as the study trials whereas Lures were presented for 1.5s if the study counterpart was presented for 1s, and vice versa if the study trial was presented for 1.5s.

Stimuli for the task consisted of randomly sampling 324 images from the 570 stimuli used in the Mnemonic Similarity Task (Stark et al., 2019) stimuli sets C-E, type “a”, for each participant. The 324 stimuli were then assigned as either Target (120), Lure (120), or Foil (84) trials. Targets and Lures repeated after either 4 or 12 stimuli (Lag = 4, 12). With 324 stimuli, 240 of which repeated once, the total number of trials was 564. Duration and Lag were counterbalanced such that (a) an equal number of stimuli were presented for both durations(282 each of 1 and 1.5s), (b) an equal number of Targets were presented for both durations (60 1s, 60 1.5s), (c) an equal number of Lures increased and decreased in stimulus duration (60 increase, 60 decrease), (d) an equal number of Foils were presented for both durations (42 each), and (e) an equal number of Targets and Lures repeated at Lag = 4 or 12 (60 Lag = 4, 60 Lag = 12). Further, each Lag (e.g., 4) had an equal number of trial durations for Targets (30 1s, 30 1.5s) and Lures (30 increase, 30 decrease). The stimuli were presented in 6 blocks of 94 trials, where each block was counterbalanced for stimulus type (Target, Lure, Foil), duration, and lag. Stimuli were presented in a pseudo-random order with the constraints of Lag and number of trials. Each block contained 20 Targets (5 for each combination of Lag and Duration), 20 Lures (5 for each combination of Lag and In/Decrease), and 14 Foils (7 of each Duration). The number of trials was selected in order to investigate potential interactions of Lag and Increase or Decrease, and further, to yield sufficient trials for modeling the hemodynamic response function in a subsequent experiment (see below).

### 2.3 Statistical Analyses

An adjusted signal-detection naming convention was adopted (Table 1) as there were two options to incorrectly identify Targets and Lures in this experiment. A correct identification of Targets was termed “Hit-Same” (Hit_Sa_) and incorrect identification was called either “Miss-Long” (Miss_L_) or “Miss-Short” (Miss_S_) according to whether the participant indicated if the Target duration increased or decreased with respect to the study trial. For Lures which decreased in duration (study trial = 1.5s, test trial = 1s), correctly identifying the stimulus was termed “Correct Rejection-Short” (CR_S_), and incorrectly identifying the stimulus was termed either “False Alarm-Long” or “False Alarm-Same” (FA_L_, FA_Sa_), according to whether the participant indicated that the Lure was presented for a longer or same duration as the study trial, respectively. For Lures which increase in duration (study trial = 1s, test trial = 1.5s), correct identification was called “Correct Rejection-Long” (CR_L_). Incorrect identification of the increasing Lure was termed either FA_Sa_ or “False Alarm-Short” (FA_S_). Note that while some response types received the same classification, such as “Hit_Sa_” resulting from correctly identifying a Target of either 1 or 1.5s, individual classifications were maintained separate during analyses.

**Table 1:** Response behavior classifications. Stimuli were presented for either 1 or 1.5s. Test trials were either Targets or Lures, where a Lure stimulus duration differed from the study trial by 0.5s. “Lure, decrease (1s)” corresponds to a study trial of 1.5s, and “Lure, increase (1.5s)” corresponds to a study trial of 1s. Participants responded whether the test trial was presented for Longer (L), Same Time (Sa), or Shorter (S) than the study trial.

| Stimulus | Response |  |  |
| --- | --- | --- | --- |
|  | Longer | Same Time | Shorter |
| Target (1s) | Miss$_L$ | Hit$_{Sa}$ | Miss$_S$ |
| Target (1.5s) | Miss$_L$ | Hit$_{Sa}$ | Miss$_S$ |
| Lure, Decrease (1s) | $FA_L$ | $FA_{Sa}$ | $CR_S$ |
| Lure, Increase (1.5s) | $CR_L$ | $FA_{Sa}$ | $FA_S$ |

A separate d′ calculation was conducted for the 1 second and 1.5 second Target durations in order to identify the response criteria, according to the adjusted formulas

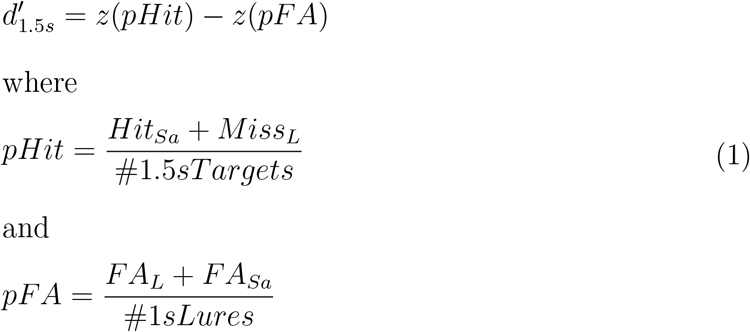

Note that while Hit_Sa_ and FA_Sa_ appear in both formulas, these response bins are kept separate in response to their respective stimulus duration.

D-prime scores were then used as the dependent variable to investigate whether or not a Lag, Duration, or interaction effect exists using a two-factor within-subject ANOVA utilizing a Greenhouse-Geisser correction for any violations of sphericity, and d′ scores were individually tested against zero using the Student’s t to determine if participant behavior differed from chance. A d′ score significantly different from 0 would indicate that participants are sensitive to changes of stimulus duration on the order of 0.5 seconds, testing hypothesis(a). A test of whether the d′ for stimuli of Lag=4 that is significantly greater than the d′ associated with Lag=12 tested hypothesis (b), and finally, a test of whether d′ scores are greater for decreases in duration relative to increases tested hypothesis (c). The multiple comparisons correction method “FDR” was utilized where appropriate.

## 3 Results - Behavioral Experiment 1

Four participants were excluded due to computer malfunction, leaving a total n=36. Outliers were detected according to the 1.5 *×* IQR method; replacing outlier values with the median value had no effect on the significance of test statistics and so all statistics reported reflect the data sets where outliers were included. A two-factor within-subject ANOVA failed to detect any effect of Duration, Lag, or an interaction thereof on d′ scores (all p-values *>* 0.05), while one sample t-testing revealed that 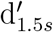 and 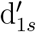 scores differed significantly from zero (*t*(34) = 14.07, *p*_*F DR*_ *<* .001, 95% CI[1.13, 1.52]; *t*(34) = 13.83, *p*_*F DR*_ *<* .001, 95% CI[1.21, 1.63], respectively). Further, these d′ scores did not differ from each other according to paired t-testing (*p*_*F DR*_ *>* .05; Figure 2, Experiment 1). Together, the evidence suggests that participant performance differs significantly from chance and that participants are equally sensitive regardless of whether the Lure increases, decreases, or appears after Lag 4 or 12.

**Figure 2:**
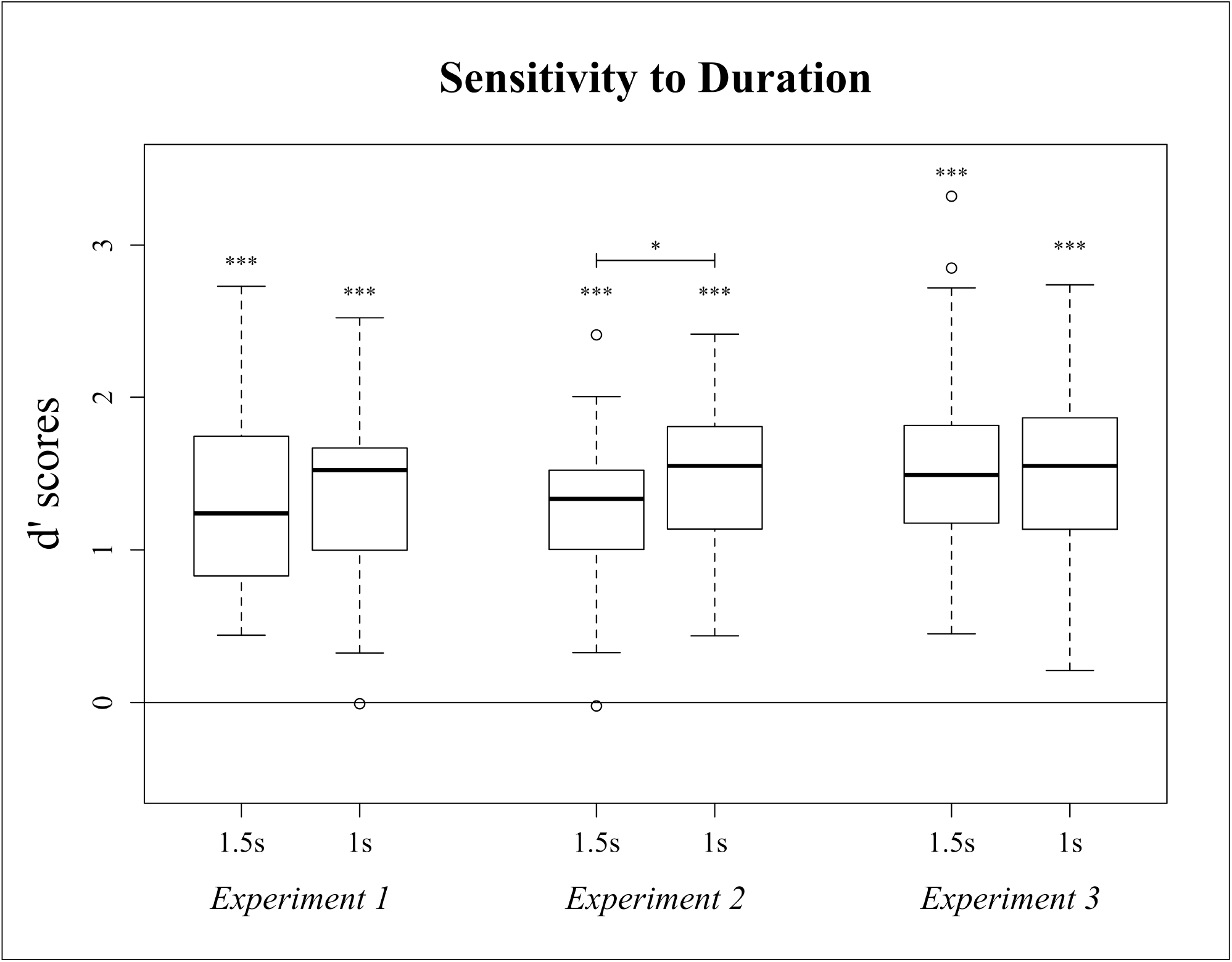
Boxplot of d′ scores. 1.5s = 1.5 second Target, 1 second Lure; 1s = 1 second Target, 1.5 second Lure. *** = p *<* .001, * = p *<* .05.

## 4 Methods - Behavioral Experiment 2

The previous task was designed with sufficient trials to investigate potential interactions of Duration and Lag as well as produce a sufficient number of trials, when subdivided by behavior, to model a hemodynamic response function. As no effect of duration, lag, or an interaction thereof was detected in the previous analyses, it was possible for us to reduce the number of trials in order to (a) decrease participant burden and (b) decrease scan cost while maintaining the necessary number of trials for MRI analyses. To this end, the number of trials in each category was reduced by exactly one-half, yielding 60 Targets, 60 Lures, and 42 Foils. In order to ensure that such a reduction would not affect the behavioral outcome, 34 new participants (16 Female, Age = 20.1 *±*1.7) were recruited to participate in the shortened version of the experiment. Participants gave written informed consent prior to participation. All methodologies and analyses conducted, save the reduction of trial items, remain identical to those reported above. Participants were compensated for their time via course credit.

## 5 Results - Behavioral Experiment 2

In the reduced experiment replacing outlier values with the group median again had no effect on test statistics and so all statistics reported include outliers. One-sample t-testing on the 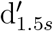 and 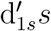 scores demonstrated that participants performed above chance (*t*(33) = 14.85, *p*_*FDR*_ *<* .001, 95%CI[1.08, 1.43]; *t*(33) = 16.44, *p*_*FDR*_ *<* .001, 95%CI[1.27, 1.64], respectively), but this time we detected that participants were more sensitive in 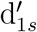 than 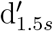 according to paired t-testing (*t*(33) = −2.22, *p*_*FDR*_ *<* .05, 95% CI[−.37, −.02]; Figure 2, Experiment 2).

## 6 Methods - fMRI Experiment

### 6.1 Participants

Forty healthy, young-adult participants were recruited from the university student population in order to investigate any potential neural correlates of discrimination for temporal duration. All participants were naïve to the task. In addition to the inclusion and exclusion criteria defined above, exclusion criteria also consisted of incompatibilities with the magnet or head coil, colorblindness and left-handedness, and the capability to remain still for an hour. The university’s Institutional Review Board approved the research, and all participants gave written informed consent prior to participation. Participants were compensated for their time via course credit, money ($20 USD), or a 3D-printed quarter-scale model of their brain.

### 6.2 Temporal Discrimination Task and Statistics

The task was identical to that in Section 4. Correspondingly, analyses of behavioral data were conducted in an identical fashion to Section 4.

### 6.3 MRI Acquisition

Functional and structural data were collected on a Siemens Tim Trio 3T MRI scanner at the Brigham Young University MRI Research Facility, using a 32-channel head coil. T1-weighted, T2-weighted, and T2*-weighted scans were collected from each participant. The T1-weighted structural scan was acquired using a magnetization-prepared rapid acquisition with gradient echo (MP-RAGE) sequence, according to the following parameters: TR = 1900 ms, TE = 2.26 ms, flip angle = 9^*°*^, FoV = 218 *×* 250, voxel size = 0.97 *×* 0.97 *×* 1 mm, slices = 176 interleaved. A high-resolution T2-weighted scan was used to image hippocampal subfields according to the following parameters: TR = 8020 ms, TE = 80 ms, flip angle = 150^*°*^, FoV = 150 *×* 150 mm, voxel size = 0.4 *×* 0.4 *×* 2 mm, slices = 30 interleaved; the T2 data were acquired perpendicular to the long axis of the hippocampus. High-resolution multi-band echo-planar scans were collected from each participant according to the following parameters: Multi-band factor = 4, TR = 2200 ms, TE = 45.2 ms, flip angle = 90^*°*^, acquisition matrix = 100 *×* 100 mm, voxel size = 1.8 mm^3^, slices = 72 interleaved; the first 11 TRs were discarded to allow for T1 equilibration. Finally, a reverse blip echo-planar scan was collected using the same parameters as the previous protocol save that direction of acquisition was opposite (P*>>*A) and only 10 volumes were collected.

### 6.4 MRI Data Pre-Processing

All pre-processing was conducted using the software packages dcm2niix (Li et al., 2016), Analysis of Functional NeuroImages (AFNI; Cox, 1996), convert 3D (c3d; Yushkevich et al., 2006) and Automatic Segmentation of Hippocampal Subfields (ASHS; Yushkevich et al., 2015). AFNI and Mango (Lancaster et al., 2012) were used for visualizations.

T2*-weighted DICOMs were converted into NIfTI files. Field distortion was then corrected by incorporating information from the reverse blip scan. Briefly, the median value of all voxels from concatenated blip and experiment functional scans were individually calculated and then the nonlinear transformation between the blip and experimental files were computed thereby producing warping and reverse-warping vectors. These reverse-warping files were then used to “unwarp” the experimental files, recovering some signal in regions of high signal distortion. The experimental volume with the minimum number of voxel outliers was then extracted to use as a volume registration base. To move data into template (MNI 152) space, the calculations of (a) rigidly aligning the structural to the registration base (using an lpc+ZZ cost function), the (b) non-linear warp of the structural file in original space to template space, and (c) the volume registration (rigid transformation) of each volume to the registration base were concatenated and applied to the functional data in native space such that all volumes were aligned with the template utilizing a diffeomorphic transformation via a single interpolation. A binary extension mask corresponding to regions within the domain with sufficient minimum signal was constructed; data within this region were scaled by the mean signal.

Single-subject regression used a generalized-least-squares fit utilizing a nonlinear residual maximum likelihood estimate to determine the best fitting autoregressive-moving-average model, where the time series corresponding to an eroded white matter mask was utilized as a nuisance regressor. Centered motion regressors for six degrees of freedom were included, and volumes that were (a) outliers in terms of total volume signal (*>*10% of the voxels in the volume had outlier signal) and (b) either preceded or were involved in a motion event (*>* 0.3^*°*^ rotation) were censored. Regressors for the various behaviors (described below) were modeled with a duration modulated block function. Two single-subject regressions were conducted, where the BOLD signal associated with behavioral responses was modeled using a boxcar convolution with the canonical hemodynamic response. First, test trials (when participants made their response) were modeled for correct and incorrect responses to Targets (Hits and Misses, respectively) and Lures (Correct Rejections and False Alarms, respectively). The hemodynamic signal associated with seeing the test trial stimulus was not modeled, as the aim of this study was to investigate processes associated with discriminating between two differing pieces of temporal information. Instead, the time between the termination of the test stimulus (when the participant had the relevant temporal information) and the memory decision was modeled. Second, the encoding of the stimuli (when the stimulus was onscreen) were modeled for subsequent behavioral responses; the entire stimulus duration was modeled. More specifically, the encoding single-subject regression modeled the hemodynamic response of the participants during the study trial of the stimulus, and this encoding session was coded for subsequent performance on the same stimulus. That is, the hemodynamic responses during the encoding of the stimuli which resulted in subsequent Hits were modeled separately from the encoding of the stimuli which resulted in subsequent Misses, FAs, and CRs. Finally, participants that had more than 10% of their total volumes censored would have been excluded from all subsequent group-level analyses but no participant reach this criterion. Data were blurred using a full-width half-maximum Gaussian kernel of a size approximately 2 times larger than the T2*-weighted voxel.

For a priori analyses, behavioral parameter estimates (*β*-coefficients) were extracted from anterior and posterior hippocampal regions of interests (ROIs) from each deconvolution utilizing masks that were previously generated in template space. Briefly, ASHS was used to segment the medial temporal regions of high-resolution T2-weighted scans for a number of participants. Joint Label Fusion (H. Wang et al., 2013) then utilized this set of segmentations to generate template hippocampal subfield priors. For this experiment, template MTL masks were split along the anterior-posterior axis (Thavabalasingam et al., 2019) at X=-20, the CA2, CA3, and DG masks were combined, and then masks were resampled into functional resolution. Any voxels labeled by more than one mask were excluded, thereby producing anterior and posterior, right and left Subiculum, CA1, and CA2/3/DG masks. A two-factor within-subject MANOVA was then used to investigate the role of hippocampal structures in the modeled behaviors, where factor A was the ROI, factor B the behavioral responses, and the parameter estimate was the dependent variable. ROIs in the left hemisphere were treated as independent from the right, and a separate analysis was conducted on the anterior and posterior masks for both the encoding and response estimates. The FDR method was used to correct for multiple comparisons.

For exploratory analyses, an intersection mask was constructed, which indicated where meaningful signal exists for both T1- and T2*-weighted scans across all participants, and multiplied with a gray matter mask that had been generated in template space, thereby producing an intersection-gray-matter mask that was used to constrain the number of voxels on which analyses were conducted. Further, a small volume mask was generated in a similar fashion but only using masks associated with the gray matter of bilateral hippocampi, entorhinal cortex, perirhinal cortex, and parahippocampal cortex. Noise was modeled by calculating the autocorrelation function of single-subject regression residuals from voxels which lay within the generated masks, and these parameter estimates were then used in Monte Carlo simulations to generate multiple-comparison correction criteria. A multivariate model was used to investigate any regions that show differential signal associated with the various behaviors modeled where the behaviors were modeled as within-subject factors. Clusters which showed differential activity were then used to extract mean parameter estimates in order to report the (a) difference in parameter estimates and (b) statistics. All scripts used for running these experiments and analyzing the data are available at https://github.com/nmuncy/MST_Temporal, and data are available at the DOI 10.18112/openneuro.ds002655.v1.0.1.

## 7 Results - fMRI Experiment

### 7.1 Behavioral Results

From the 40 participants recruited, five were excluded from analyses due to equipment problems and incompatibilities (n=3), responding with the wrong button presses (n=1), and “excessive eye watering” during the task which was reported to be “very distracting” and resulted in at-chance performance (n=1). The group size reported below is therefore n=35, 19 female, age=22.9 *±*2.7.

As above, statistics were conducted both with and without outliers. Replacing outlier values with median values had no effect on test statistics and so all analyses below include outliers. Performance was very similar to the above experiments (Figure 2, Experiment 3). Participants d′ scores differed significantly from 0 according to one-sample t-testing for both increases and decreases in duration (*t*(34) = 13.9, *p*_*FDR*_ *<* .001, 95% CI[1.35, 1.81]; *t*(34) = 16.4, *p*_*FDR*_ *<* .001, 95% CI[1.32, 1.69], respectively). In addition, and similarly to the results of Experiment 1 but not Experiment 2, no difference was detected between sensitivity to duration increases versus decreases according to two-sample t-testing (*t*(34) = 0.65, *p*_*FDR*_ = .52, 95% CI[−.16, .31]).

### 7.2 FMRI Results - Encoding

As no effect of Lag or Duration was detected in previous analyses (Section 3) and participants performed equally well regardless of increasing or decreasing the Lure duration (Section 7.1), responses were collapsed across both Lag and Duration. Additionally, types of Misses, Correct Rejections, and False Alarms (Table 1) were collapsed across as well. Such collapsing resulted in average of 28 Hits, 31 Misses, 31 Correct Rejections, and 23 False Alarms.

The effect of encoding on task performance was assessed by investigating the BOLD response to stimulus presentation that preceded a task response (Hit, CR, etc.). For the a priori ROIs, no significant effect was detected via a two-factor within-subject MANOVA for either ROI, Behavior, or an interaction thereof (all p-values *>* .05 once correcting for sphericity via the Greenhouse-Geisser method) for both the anterior and posterior regions of the hippocampus.

An small-volume corrected analysis of the MTL, using a mask corresponding to hippocampus, entorhinal, and parahippocampal cortex, revealed two clusters with differential activity during study trials that preceded a test response when using the thresholding criteria of k = 6, NN = 1, two-sided comparison, p *<* .001, FWE = .05. Neuroanatomic classification of the clusters for these and subsequent analyses were determined by the criteria established by Frankó et al. (2014). The left entorhinal cortex was more active during encoding trials preceding a Hit versus a Miss (LEC, Figure 3, Table 2) while the right hippocampus was more active during encoding trials preceding a FA than a CR (RHC, Figure 3 Middle, Table 2). Finally, an exploratory whole-brain analysis failed to detect any clusters which showed differential signal when using the thresholding criteria of cluster size k = 20, nearest neighbor (NN) = 1, two-sided comparison, p *<* .001, FWE = .05.

**Table 2:**
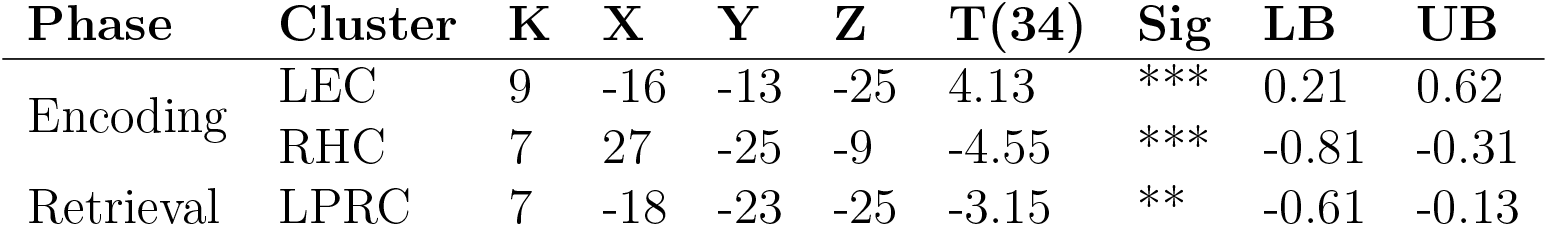
Cluster descriptions of small-volume corrected analyses. K = cluster size, X-Z = peak MNI coordinate. T(34) = paired t-test with 34 df, LB UB = lower and upper bound of 95% confidence interval. LEC = left entorhinal cortex, RHC = right hippocampus, LPRC = left perirhinal cortex. *** = p *<* .001, ** = p *<* .01.

| Phase | Cluster | K | X | Y | Z | T(34) | Sig | LB | UB |
| --- | --- | --- | --- | --- | --- | --- | --- | --- | --- |
| Encoding | LEC | 9 | -16 | -13 | -25 | 4.13 | *** | 0.21 | 0.62 |
|  | RHC | 7 | 27 | -25 | -9 | -4.55 | *** | -0.81 | -0.31 |
| Retrieval | LPRC | 7 | -18 | -23 | -25 | -3.15 | ** | -0.61 | -0.13 |

**Figure 3:**
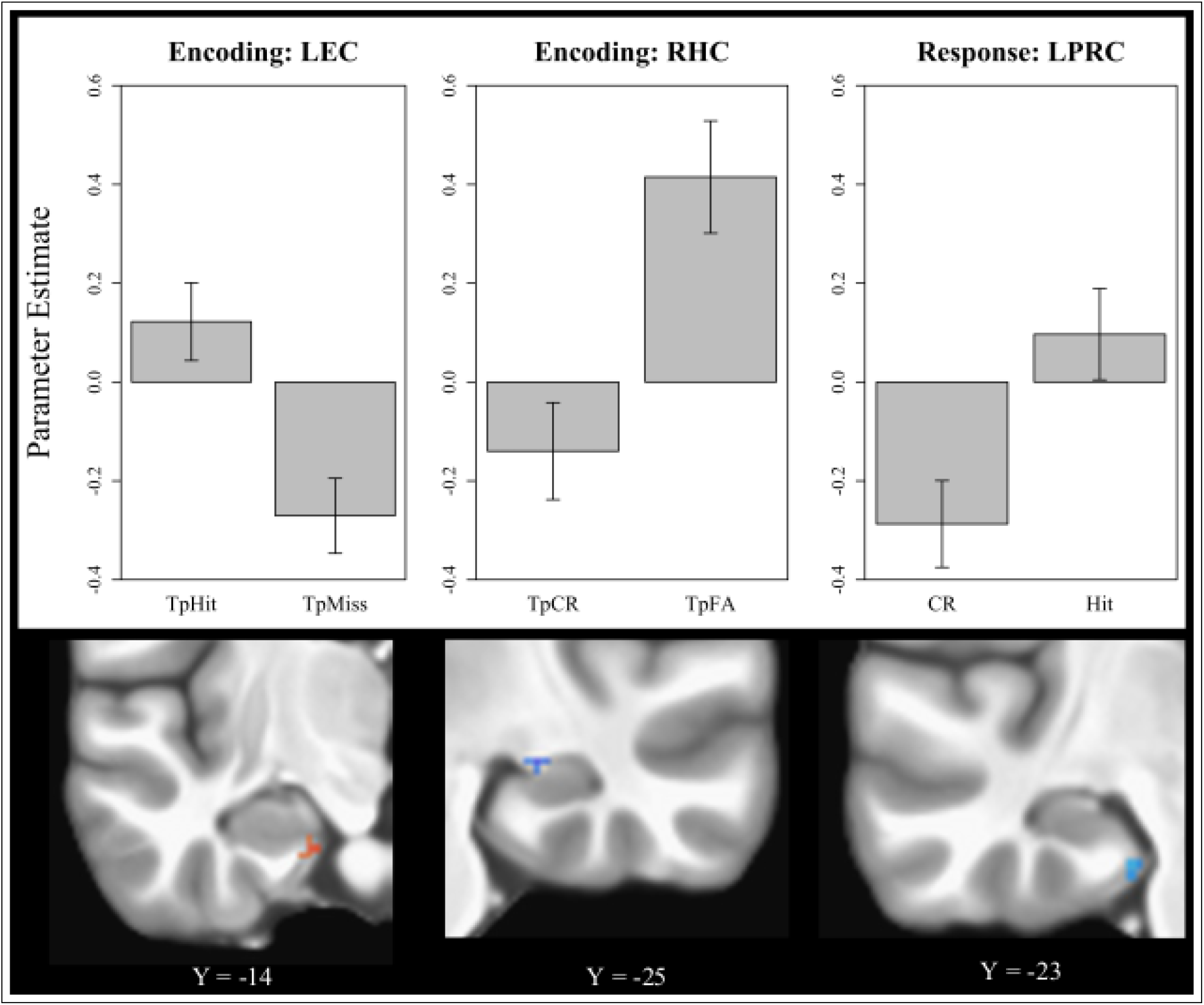
Regions engaged in duration change detection. Left and Middle: left entorhinal cortex (LEC) and right hippocampus (RHC) are differentially engaged during encoding trials preceding task performance. Right: Left perirhinal cortex (LPRC) is differentially involved in successful test trials. “Tp” = trial preceding a test response.

### 7.3 FMRI Results - Retrieval

To investigate processes associated with task performance the hemodynamic response function was modeled for the time in which participants responded to Target and Lure stimuli. A priori investigations into the subregions of the medial temporal lobe failed to detect a significant interaction between region and behavior in either portion of the hippocampus. A small-volume corrected voxel-wise analysis, however, detected a cluster in the left perirhinal cortex that was differentially engaged in correct responses on test trials (LPRC, Figure 3 Right, Table 2). That is, this cluster was more active during the correct identification of Targets (Hits) than during the correct identification of Lures (CRs). Finally, an exploratory whole-brain analysis using the same thresholding criteria as was reported above failed to detect any clusters that demonstrated differential activity for the various task behaviors.

## 8 Discussion

This set of experiments investigated whether mnemonic discrimination was possible for stimulus duration and identified functional neuroanatomic correlates thereof. The task (Figure 1) involved serially presenting images of everyday items in a continuous-recognition paradigm for either 1 or 1.5 seconds. Following the study trial of a stimulus the participants performed an orienting task wherein they made decisions from memory about physical properties of the stimulus. If a stimulus repeated, it was either presented for the same duration (Targets) or for a difference of 0.5 seconds (Lures); in these cases, the participants were explicitly instructed to attend the presentation duration and subsequently tasked with deciding whether a change of stimulus duration occurred. Such a small change in stimulus duration was intentionally selected in order to elicit a sufficient number of Misses and FAs for functional BOLD analyses. The paradigm was conducted three times in naïve groups of participants, Experiments 2 and 3 serving to replicate the behavioral findings of Experiment 1 and bringing the total number of participants that performed the task to n=69.

These experiments had several findings. First, behavioral evidence indicated that participants were successful at remembering and detecting changes of stimulus duration on the order of 0.5 seconds and also that they were equally sensitive regardless of the direction of change (Section 3). Importantly, these findings appear to be robust, as the d′ scores were stable across interenal replications with three independent groups of participants. With respect to these behavioral findings, we reject the null of hypothesis (a), given that participants had d′ scores which differed significantly from zero, but fail to reject the null of hypotheses (b) and (c); participants were equally sensitive at Lag 4 as Lag 12, as were they to increases and decreases in duration.

These findings of sensitivity to timing changes of less than one second replicate those of both Barnett et al. (2014) and Palombo et al. (2019), where both sets of experiments indicated that the human memory system is sensitive to second and sub-second duration changes. This experiment differs from that of Barnett et al. (2014) and/or Palombo et al. (2019) in that we (a) had participants detect temporal changes on a trial-by-trial basis rather than for an entire set of stimuli, (b) indicate the direction of suspected temporal change rather than using a match-mismatch paradigm, and (c) that this task set participants to work within long-term, episodic memory rather than short-term memory. Despite the differences, our findings are quite similar to Palombo et al. (2019) who noted, as we do here, that participants had strong d′ scores and were sensitive to both increases and decreases in duration. Somewhat differently, test trials in this experiment were likely presented within the windows of both short- and long-term memory, when using a reference point of approximately 30 seconds; Lag 4 test trials occurred between 17.2 and 19.2 seconds after the study trial and Lag 12 test trials would occur between 51.6 and 57.6 seconds after their respective study trial. It is interesting, then, that participant performance on both Lags was equal. We had anticipated higher d′ values on Lag 4, values similar to those presented by Palombo et al. (2019), than on Lag 12. From our data, however, it appears that the same discriminating processes are supporting both Lag 4 and 12 performance, even though these trials may be classified as working and episodic memory, respectively. This may potentially give insight into why the MTL patients of Palombo et al. (2019) were successful at temporal discrimination when working within the time frame of less than 30 seconds, given that working memory processes are not necessarily MTL dependent. Another possible interpretation of the findings presented here is that the task was sufficiently difficult such that it overwhelmed a limited short-term memory, and any subsequent test performance was then due to episodic memory discrimination.

Second, two clusters showed differential activation during the encoding which was predictive of subsequent test performance. A cluster in the right hippocampus (Figure 3, middle) was differentially engaged in encoding such that the cluster was more active preceding the incorrect detection of Lures (FA) compared to a correct detection (CR). This pattern of activity is consistent with the deleterious effect that the default mode network has on subsequent test performance (Kim, 2011). It should be noted, though, that a second, a posteriori whole-brain analysis did not identify any additional clusters to those detected in the small-volume corrected analysis. As such, the lack of additional default-mode findings substantially weakens the interpretation of this hippocampal cluster. A second interpretation, then, is that this cluster is indicative of inefficient encoding such that subsequent memory testing elicits mnemonic generalization, i.e., subsequent forgetting.

Next, the left medial entorhinal cortex (mEC) demonstrated differential activity which predicted subsequent Target performance, in that the cluster was more active preceding a Hit relative to preceding a Miss (Figure 3, left). This cluster appears to be critically involved in encoding the temporal properties of the stimuli, as visual information remained identical between study and test, a finding that is consistent with the previous literature: numerous studies have detected cells with a selectivity for temporal information within the mEC. For instance, Kraus et al. (2015) demonstrated mEC grid cell activity which signaled for elapsed time in addition to distance traveled, showing an integration of time and distance, and Heys and Dombeck (2018) demonstrated both that while the mEC is sensitive to elapsed time, it can also remap its sensitivity according to contextual task demands (also, see Sugar and Moser, 2019). Robinson et al. (2017) showed that the mEC helps support the hippocampal CA1 representation of time, and further, Suh et al. (2011) demonstrated that the monosynaptic side of the perforant pathway, the connection between the mEC and CA1, is integral for temporal associations. This hippocampal structure, the CA1, is downstream from the mEC and has been strongly implicated in the temporal aspects of memory (Allen et al., 2016; Cai et al., 2016; Dimsdale-Zucker et al., 2018; MacDonald et al., 2011; Mankin et al., 2012; Salz et al., 2016; Suh et al., 2011; Y. Wang et al., 2015). Finally, inactivation of the mEC is also associated with disruptions in temporal processing (Heys & Dombeck, 2018; Robinson et al., 2017; Sugar & Moser, 2019)). As such, the mEC is proposed to bind information of different modalities (i.e., item, spatial, temporal) as well as providing a scaffold of spatiotemporal information to the hippocampus (Sugar & Moser, 2019).

Further, Maass et al.(2015) and Schröder et al. (2015) demonstrate that the mediolateral divisions of the rodent entorhinal cortex do not translate to the human entorhinal cortex, but that the human entorhinal cortex is functionally divided along an anterolateral-posteromedial axis such that the human homologue of the rodent mEC is the posteromedial entorhinal cortex (pmEC). Maass et al. (2015) shows the PHC is functionally coupled with the pmEC, implicating this region in processing contextual information as opposed to item information which tends to be sent from the PRC to the lateral anterior EC (Aminoff et al., 2007; Diana et al., 2013; Hsieh et al., 2014; Jenkins & Ranganath, 2010; Naya et al., 2017; F. Wang & Diana, 2017). Critically, the cluster detected in this experiment during the encoding phase of the experiment entirely corresponds with the pmEC. Further still, Schröder et al. (2015) demonstrated that pmEC forms part of the posterior-medial system, a system which carries spatial information. The rodent and human literature in conjunction therefore implicate the pmEC in processing not just spatial but contextual information which varies in a dimension (i.e. temporal information) according to task demands, a notion that is reminiscent of Salz et al. (2016) who noted that temporal processing in the hippocampus mirrors that of spatial processing. In the current experiment, then, the detected pmEC cluster is believed to be processing the temporal context information associated with each item.

Finally, a cluster within the left perirhinal cortex (PRC) was differentially engaged during the test phase such that it was more active during Hits than Correct Rejections (Figure 3, right). Similarly, recent work implicated the PRC in processing temporal information during the test phase of an experiment: Montchal et al. (2019) had participants first view a television episode and then tested participants by having them place still-frames from the episode on a timeline, a task designed to test temporal precision memory. While the paper emphasized the roles of the anterior-lateral EC and subregions of the HC, Montchal et al. (2019) nevertheless demonstrated that the PRC was differentially engaged in high versus low accuracy for temporal information. In a separate but related vein, the Binding of Item and Context theory (Diana et al., 2013) has proposed a functional dissociation in the medial temporal lobe where the parahippocampus processes contextual information, the PRC item information, and the hippocampus serving to bind together item and context; the role of the PRC in such a model has recently been supported (Diana et al., 2013; Hsieh et al., 2014; F. Wang & Diana, 2017). Together then, the PRC has been implicated in processing both temporal and item information. Such a conjunction of item and time is supported by F. Wang et al. (2017), who noted PRC activation associated with the retrieval of contextual temporal information, perhaps due to the ‘unitization’ of the item and context (Ford et al., 2010; Giovanello et al., 2006). More explicitly, work using single-unit recordings in non-human primates demonstrated that PRC cells fired for both item and temporal information (Naya et al., 2017). Thus, the PRC activation detected in this experiment is consistent with the conjunction of item-temporal information processing, in which the PRC differentially aides in the successful retrieval of item and temporal duration information. Indeed, increased signal from this region is associated with Hits, i.e., correctly retrieving the duration information (as the visual information was identical), while suppressed signal is associated with a mismatch detection between the Lure duration and that of the original.

A number of limitations exist for this experiment. First, while only Targets (same duration) and Lures (different duration) were tested in this experiment, participants had three response options necessitating an adjustment to the d′ calculations. This adjustment resulted in collapsing across some response types such that these adjusted-d′ scores do not represent a bias-corrected deflection point but rather indicate sensitivity to whether a change occurred. Indeed, in calculating 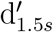 (when the Target was 1.5s and the Lure was 1s), *pHit* combined both responses that did not involve a “shorter” judgment for Targets, and *pFA* did the same. With such a calculation, a higher adjusted-d′ score therefore indicates that a participant is less likely to respond “shorter” on longer Target trials and more likely to respond the same on Lure trials, a response pattern that is related to, but separate from, accuracy in judging stimulus duration in episodic memory.

Second, and related, several response categories were collapsed across in the functional analyses: as no main effect of Lag or Duration was detected, which could have been used to justify a investigation into fMRI differences associated with these effects, the same responses from different stimulus types (such as Hit_Sa_ for both a 1.5s and 1s Target for both Lags) were combined into the same bin. This method ensured a sufficient number of trials per behavioral outcome existed such that the shape of the hemodynamic response function could reasonably be modeled, but it nevertheless may have contributed noise to the analyses. If certain regions of tissue were involved in only detecting increases, decreases, or no change in stimulus duration then such a combination would likely have introduced unnecessary noise to any existing signal. As such, our functional findings do not illustrate signal associated with correctly identifying increases or decreases of duration in Targets and Lures, but rather demonstrate fMRI signal associated with detecting temporal change between the study and test stimuli.

Third, a recent meta-analysis of the neural correlates of time in human participants (Nani et al., 2019) implicated insular, opercular, dorsomedial prefrontal, parietal, and striatal regions in sub–and supra–second non-motor time perception; we note that our study surprisingly did not identify similar regions. A number of explanations are possible. First, our experiment was designed to detect changes in fMRI activation associated with episodic memory precision for temporal duration, rather than time production, processing, or perception as was the aim of the studies included in the meta-analysis. Second, we note that all studies included in the meta-analysis were published prior to the Eklund et al. (2016), save for three Alhaj et al., 2016; Apaydın et al., 2018; Üstün et al., 2017. Our MRI analysis methods include updated steps to better deal with the multiple comparisons issue, utilizing methods developed in response to Eklund et al. (2016). These methods are more conservative than those employed by the studies included in the Nani et al. (2019) meta-analysis. Additionally, we had a larger number of participants than all studies included in the meta-analysis, save one (Tomasi et al., 2015), which helps to guard against the inflated Type-1 (or false positive) error and effect sizes that result from smaller sample sizes (Button et al., 2013; Friston, 2012). Compounded with these differences in sample size and analytic methods, the voxel detection method employed in this analysis, 3dMVM and using criteria informed by modeling the ACF via 3dFWHMx, will only detect a portion of significant voxels in any analysis, missing both larger voxel clusters with “lower” significance and smaller clusters with “higher” significance (Cox, 2019), due to the requirement of supplying a minimal cluster size and maximal statistic threshold. Together, then, we speculate that our failure to detect similar findings is due to (a) differences in cognitive processes (episodic memory precision versus time perception, processing, etc.) as well as (b) a highly conservative analytic approach that minimized our experiment’s susceptibility to an inflated effect size and Type-1 error rate. On this latter point, it is possible that the lack of results is also partially due to Type-2 (or false negative) errors, given the limitations of cluster detection by 3dMVM (Cox, 2019).

Fourth, while the hippocampal subregion CA1 has been shown to be critically involved in temporal processing for memory tasks (Allen et al., 2016; Cai et al., 2016; Dimsdale-Zucker et al., 2018; MacDonald et al., 2011; Mankin et al., 2012; Montchal et al., 2019; Robinson et al., 2017; Salz et al., 2016; Y. Wang et al., 2015), our results did not replicate these findings. This is likely due to the fact that the rodent studies which have implicated the CA1 have largely involved requiring the rodent to maintain information over time (e.g., Kraus et al., 2015; Pastalkova et al., 2008), information which is not necessarily temporal and is commonly a previous decision. Our experiment instead involved the retrieval of temporal duration information, and likewise it is unsurprising that we detected signal associated with tissue in a different region. With respect to previous studies involving humans (Aminoff et al., 2007; Clewett & Davachi, 2017; Diana et al., 2013; El-Kalliny et al., 2019; Hsieh et al., 2014; Jenkins & Ranganath, 2010, 2016; Montchal et al., 2019; F. Wang & Diana, 2017), our results are similar in nature and any differences are likely due to the fact we here tested for episodic memory precision of temporal duration rather than temporal order or sequence, or working memory for temporal duration (Palombo et al., 2019). Of particular note, Thavabalasingam et al. 2019 recently demonstrated that the anterior hippocampus, and more specifically the right anterior CA1, contained a pattern of activity that seemingly represented the conjunction of image and time as tested by a sequence decoding classifier (and impressively that this pattern of activity was present both during recognition as well as mental replay). When investigating image and time separately, however, a temporal decoding classifier failed to detect any pattern of activity in either the anterior or posterior hippocampus. Given that we tested for episodic memory precision for temporal duration divorced from item or sequence, our lack of CA1 and hippocampal findings mirror those of Thavabalasingam et al. 2019. We posit that the findings of Thavabalasingam et al. 2019 supports the Binding of Item and Context theory (Diana et al., 2013), where hippocampal CA1 is sensitive to a conjunction of context and item, and that our lack of CA1 findings is the result of targeting temporal context alone.

Together, then, we partially reject hypothesis (d), as we did find differential activation in the MTL associated with the task, but not in the hypothesized region of CA1. Further, we reject hypothesis (e), as no differential signal was detected in the anterior versus posterior HC. It is possible that we failed to detect differential signal in these regions due to collapsing across too much variance, or that the selected contrasts did not reflect differences in the signal, or perhaps even due to the explicit nature of our task. Either way, one must take caution in interpreting null results (Chen et al., 2017) and note only that we were unable to reject the null of these hypotheses.

In summary, we developed a novel paradigm where we target episodic memory precision for time by presenting participants with stimuli of different duration and tasked them with detecting a change in duration. We found that participants were sensitive on the order of 0.5 seconds, a finding which was demonstrated in three different groups of participants. We also demonstrate that the left posteromedial entorhinal cortex is differentially involved during the encoding of items such that greater activity in this region is predictive of subsequent Hit versus subsequent Miss. Finally, the left perirhinal cortex was noted to be differentially engaged during successful test performance, being more active during Hits than Correct Rejections, activation which is consistent with previous work demonstrating that this region is sensitive to the conjunction of item and temporal context.

## Acknowledgments

The authors have declared that no competing interests exist.

